# Decipher cell communication with attention: CLARA

**DOI:** 10.1101/2025.03.09.642280

**Authors:** Honglin Wang, Yeonsoo Chung, Dong-guk Shin

## Abstract

Cell-cell communication is crucial in maintaining cellular homeostasis, cell survival and various regulatory relationships among interacting cells. Thanks to recent advances in spatial transcriptomics technologies, we can now explore how cells’ proximal information can be used to infer cell-cell communication. Most current studies on cell communication focus on reporting general interactions between cell types without cell proximal information. While few methods detailing communication for each cell pair, two main draw backs can be found in these methods. I. These cells has fail to provide a reasonable cell communication radius and depend on pre-define. II. These methods tend to find cell communication based on highly expression genes which are inferred by comparing global gene expression values instead of local expression value. This will lead to miss identifying local high expression values.

Here we present a cell-cell communication inference framework, called GACom, which converts a cell communication problem into a natural language process problem and infers each cell pair’s communication based on the cell-transformed word relationship with an altered attention mechanism named ligand-receptor attention. We used one mouse embryo SeqFISH dataset and one human cartilage dataset to demonstrate the performance of CLARA. In Mouse embryo dataset, we explore the communication pattern through Vascular endothelial growth factor. In human cartilage dataset, we explore how the DKK1→LRP6 and IL6→IL6ST change at normal and osteoarthritis. We also demonstrate how the CLARA can provide a new view for cell sub grouping including the cells that are sending the ligand and the cell that are receiving ligand. The biological process analysis shows the rationality of the grouping strategy. CLARA is an attention-based cell communication inference solution that leverages spatially or virtually spatially resolved single-cell transcriptomic data to predict cell-cell communication, further providing a strategic grouping approach based on cell communication insights.

## Introduction

Multicellular organisms rely on the precise coordination of cellular activities, which hinges upon effective intercellular communication among the various cell types and tissues within the organism^1–3^. Thus studying cellular functions increasingly requires understanding how cells communicate with each other given a temporal and spatial context. It is known that signaling events are frequently orchestrated through diverse protein interactions such as ligand-receptor, receptor-receptor, and extracellular matrix-receptor interactions. In response, recipient cells initiate downstream signaling via their respective receptors, typically culminating in the modulation of transcription factor activity and subsequent changes in gene expression.

Historically, cellular communication could only be studied via in vitro experiments consisting of one or two cell types and a selected number of genes. Thanks to advances in single-cell technologies and spatial transcriptomics, which measure gene expression at a single-cell resolution, scientists can now attempt to identify cell-cell communication as well as cell type communication. Currently, there are many computational tools that infer cell communication from single-cell transcriptomics. CellChat^4^ computes cell type communication probabilities using the law of mass action and considers the geometric means of ligand and receptor expressions weighted by their agonists/antagonists. CellPhoneDB^5^ generates a list of the most statistically significant ligand-receptor interactions by generating a null distribution by permuting cell cluster labels. MISTY^6^ constructs a gene communication signaling network within and between cell clusters. stLearn^7^ computes a ligand–receptor coexpression score related to cell type diversity at individual ‘spots’ to identify spatial regions with intensive signaling activity. Cell2Cell^8^ infers communication distance using a Gaussian mixture model; it uses a Bray–Curtis–like score to model interactions and optimizes Spearman correlation between distances and interaction scores. SpaOTsc^9^ and COMMOT^10^ generate a cell communication matrix for a given signaling pathway by solving an optimal transport problem. These methods, however, often summarize data at the cell group or type level, potentially overlooking the precise and individual interactions between specific cell pairs. This limitation can obscure vital insights into the cellular microenvironment, which are crucial for understanding tissue functionality and the pathogenesis of diseases.

Reporting each cell pair communication could be computationally expensive, as in biological tissue, cells are surrounded by thousands of cells and considering one more cell will increase the computational workload exponentially. With the development of natural language processing (NLP), a novel machine learning mechanism, the self-attention^11–13^ mechanism, has been developed to compute the interaction between different words in a sentence within linear time. Self-attention is a mechanism in transformer models that calculates the relevance of all words of the input data to each other, allowing the model to weigh information dynamically and contextually by computing attention scores for each element in a sequence relative to all other elements. In this research, we transform the cell communication inference question into an NLP question and developed a framework for inferring cell-cell Communication with Ligand And Receptor Attention (CLARA). There are two key innovations in the framework to enhance the understanding of cellular communication mechanisms. Firstly, we transform cell proximity information into graph representations, where cells are depicted as nodes interconnected by edges reflecting their spatial relationships. By structuring this data into “cell sentences,” we effectively capture the interactions between target cells and their neighboring counterparts within the graph framework. Secondly, we introduce a novel approach termed “ligand-receptor attention,” which is a revised version of self-attention specifically tailored for capturing ligand-receptor interactions within the constructed cell sentences. This attention mechanism operates by computing attention scores that weigh the relevance of different ligand-receptor pairs based on their expression levels and cellular context. The training targets of this framework can be either category data such as cell type, or it can be continuous data that shows the properties of the cell including biological process score. This hypothesis is supported by the premise that cells of a particular type are characterized by distinct gene expression patterns, necessitating specific ligand or receptor expression patterns for their identification^14^. Accordingly, attention scores could be interpreted as cell-cell interaction (CCI) scores, contingent upon the selection of an appropriate training target. Finally, a heuristic computational algorithm was introduced to quantify communication between neighboring cells through various attention scores based on ligand and receptor expression value in each cell pairs. Receptor-related downstream transcription factors’ expression is also taken into account in the algorithm to improve the accuracy of the inference. We showcase CLARA’s overall capabilities by applying it to both mouse embryonic Seq-FISH dataset and human cartilage single cell dataset. A systematic comparison with several existing tools for cell–cell communication is presented. And gene ontology (GO) analysis is also used to demonstrate the rationality of the new communication-based clustering strategy.

## Results

### CLARA decipher cell pairs communication score with ligand-receptor attention

CLARA is a supervised tool for extracts the latent patterns of inter-cellular communication with revised self-attention. Self-attention mechanisms offer the distinct advantage of efficiently capturing long-range dependencies in sequence data, enabling models to directly compute relationships between any two positions, regardless of their distance apart in the sequence and quickly identify the features importance based on its weight. Extending this mechanism to infer cell-cell communication enables analyzing importance of each ligand-receptor pairs in a cell pairs manner. Briefly, CLARA first leverages spatial information of cells, gene expression values, and ligand-receptor pairs to construct “cell sentences” for each cell and its neighbors (Fig.**??** A). Following this, the model employs one of the cell properties, for example cell type or biological process score information to refine the weight within the ligand-receptor attention matrix, enhancing the precision of attention scores for each ligand and receptor pair between each two cells (Fig.**??** B). The ligand-receptor attention matrix is a sparse matrix that map the ligand and receptor relationship. These attention scores, in conjunction with gene expression data, are then utilized by the cell communication inference module to deduce the communication scores among different cell pairs.

To demonstrate how CLARA infers cell communication, we applied the system through 39272 ligand-receptor-TFs triplets across 2 kinds of real world datasets including datasets from mouse embryos datasets and datasets from human cartilage. The mouse embryos SeqFish datasets^15^ has 3 separated dataset with 23 cell types, the proximity information is provided from the original source. The human cartilage sc-RNA datasets can be grouped into two type including cartilage normal(CN) and osteoarthritis(OA). There are 11039 cells in normal datasets and 10921 cells in OA^16^. Though the cell proximity information is not originally provided, multiple studies have state that human cartilage can be split into three zones including Superficial zone (S), Middle Zone (M) and Deep Zone (Z)^17–19^, based on the expression pattern of multiple marker genes. We use these genes to calculated the zonal score to stimulate the Y proximity values meanwhile, we random assign X value. (Fig.**??** A,B). Based on the zone score, we manually separate cells into three coordinate groups according to the cell density of each dataset (red line in the Fig.**??** B).

A general communication results generated from CLARA are provided in Fig.**??** C and D, we found that there 40 common cell communications from top 50 communication in all 3 mouse embryos datasets and we also notice that there are 5 unique communications ranked in top 50 communication in both normal and OA datasets. The large number of communication overlapping is expected as they are from same tissues which proves the result stability generated from CLARA.

In the following sections, we first compare the CLARA with other two methods to demonstrate the advantage of the CLARA for reducing both false positive rate and false negative rate. Then, we demonstrate to using CLARA to find the communication between different cell types through Vascular endothelial growth factor in mouse embryo. Finally, we also demonstrate the compatibility of CLARA by using continues label to find the different cell communication patterns between CN and OA.

### CLARA significant reduce the false positive and false negative rate

Cell–cell interactions (CCI) are important in all multi-cellular processes, both for normal tissue growth or maintenance, and in disease-driven change. Current methods to find biologically significant ligand-receptor (LR) interactions in any of these contexts often suffer from a common limitation, that is, high false discovery rates. For instance, scRNA-seq data lacks spatial context, meaning that interactions could be predicted between cell types that are spatially very distant from one another, and are thus unlikely to directly interact. Two strategies are applied in the current methods including: A. Using cell proximity information, as a cell rarely communicate with cells that far away. B. Using permutation test or other statistical test from global gene expression to filter out identified CCIs. However, using statistical test from global gene expression tends to produce false negative case. CLARA solves this issue by first identifying spatial neighbourhoods and form the cell sentences. This is followed by filtering out the cell communications by considering the downstream and the expression pattern within the cells and its neighbor cells within a cell community locally. CLARA solve the issues by first first identifying spatial neighbourhoods of central cell with its neighbor cells to built the “cell sentence” and find the cell communications within the cell sentence. This can significantly reduce the false positive inference. This is then followed by filtering with heuristics rule to filter it with the ligand and receptor expression relationship along with downstream Transcription factor expression pattern with in locally. As the same gene expression can vary from cell sentences to cell sentences, considering the communication local will reduce the false negative rate. With applying these two strategies, CLARA can make specific inferences about of cell interactions at both the level of individual cells or cell types with reduce false positive and false negative simultaneously.

Comprehensive bench-marking with CellChat and stLearn showed that CLARA markedly reduces false positive predictions and false negative. CellChat summaries multiple pairwise cell communication with single cell RNA-seq while stLearn is a spatial constrained cell communication inference tool. To test and/or validate this premise, we first compared CLARA performance against these two other CCI methods using a simulated dataset. The simulation established realistic gene expression values based on either scRNA-seq or spatial data, also hypothetically arranging cellular neighbourhoods as state in stlearn (**??**. A). In the comparison, CLARA and stlearn is set the same communication distance constrain. Compared with the ground-truth for cell–cell interactions in the simulated dataset (Fig. B), CellChat produced multiple false positive, while stlearn fail to identified the communication that across different cell types (an example is give in **??**. C). CLARA was the only method able to reconstruct these interactions without any additional false positive interactions and not missing the true communications (**??**. D).There are two pars of the comparison with real world mouse embryo datasets, including: ligand-receptor communication wise (**??**. E) and cell type communication wise (**??**. F). In the ligand-recepotr communication wise, CLARA has 8 overlapping in E1 and 12 in E2 dataset while it has 1 overlapping in E3 dataset when comparing with the top 50 significant results generated from CellChat and stLearn. CLARA demonstrate more overlapping with the results from CellChat than stLearn in L-R wise comparison while CLARA demonstrate more similar with the results from stLearn than CellChat in cell-type wise analysis. To validate the correctness of found communication ligand and receptors, we performed the Gene ontology(GO)^20^ analysis with identified top 50 ligand-receptors from each methods. The cell division is not shown up in top 5 identified biological process by stLearn as expected^21^.

### CLARA can provide detailed cell cell communication

To evaluate the ability of CLARA in inferring the cell cell communication, we analysis the results from the CLARA for two different datasets. In the mouse embryo dataset, we explore the communication through Vascular endothelial growth factor in mouse embryo and in the human cartilage datasets, we explore how the acidification and wnt signaling are changed when OA happens. After identified the communication, we provide a new grouping method, ligand sender and ligand receiver group, to the cells and perform the GO analysis for each group. The analysis result prove the correctness of our inferred communication.

### Explore how the communication through Vascular endothelial growth factor in mouse embryo with CLARA

In the embryonic stage, the foundational vascular network is established through a process known as vasculogenesis, which involves the *in situ* differentiation of mesodermal cells. Despite the complexity of these angiogenic processes, the molecular mechanisms that underpin them are not yet fully understood. Nonetheless, burgeoning evidence highlights the indispensable role of vascular endothelial growth factor (VEGF) in facilitating both vasculogenesis and angiogenesis across embryonic development and into adulthood^22,23^. VEGF, a key member of the platelet-derived growth factor (PDGF)/VEGF family, serves as a specialized growth and survival factor for endothelial cells. To date, this family is known to comprise two PDGFs (A and B) and five proteins closely related to VEGF—namely, placenta growth factor (PlGF), VEGF-B, VEGF-C, and VEGF-D^24–26^, with VEGF-B being particularly akin to VEGF. VEGF-B is expressed in various tissues but is most abundant in the heart, brain, skeletal muscle, and kidney, and it predominantly binds to the VEGFR-1 (FLT-1) receptor^27,28^. By using CLARA, we delve into the mechanisms through which VEGF-B interacts with FLT-1, via both paracrine and autocrine signaling pathways, across different cellular environments. The CLARA tool facilitates the visualization of cell-cell communication scores, which are quantitatively depicted for each cell pair. As demonstrated in Fig. **??**A, the communication mediated by VEGFB and VEGFR-1 is prevalent among a majority of cells, with a significant color representation in the visual output. It is observed that while most communication events are characterized by a communication score below 0.5, indicating low interaction strength, robust communication is detected at the interface between various cell types. Among these, the top ten communication events with the highest scores across three datasets involve interactions from Forebrain/Midbrain/Hindbrain to Endothelium, Neural Mesenchymal Progenitors (NMP) to Endothelium, and Spinal Cord to Endothelium, as illustrated in Fig. **??**B. Additionally, the violin plots provided by CLARA shed light on the distribution of communication coefficients among different cell types. Specifically, endothelial cells and blood progenitor cells display higher communication coefficients as receiver cells rather than as senders, suggesting a greater propensity to receive VEGFB signals from other cell types. This is corroborated by bar chart analyses, which identify endothelial cells and hematoendothelial progenitors as the primary cell types secreting VEGFB and engaging in binding with VEGFR-1 on endothelial cells (Fig.**??**C,D).

Utilizing CLARA to amalgamate communication findings affords a novel stratification method within cellular dialogues. This study categorized endothelial cells into two cohorts based on VEGFB activity: cells that secrete VEGFB and consequently bind to VEGFR-1 within the endothelium, and those that do not engage in VEGFB secretion. While the UMAP visualization does not depict a stark segregation between these groups, discernible zones where one subgroup is more concentrated do emerge. These zones signal niches in the high-dimensional landscape characterized by VEGFB secretion homogeneity (Fig.**??**E). To underscore the utility of communication in delineating cellular subgroups, differential gene expression analyses were executed between the VEGFB-secreting and non-secreting ensembles. VEGFB itself was ranked among the top 10 differentially expressed genes across all three datasets examined. A suite of other genes, including CHCHD7, HSPE1, DCTPP1, RPS14, SSBP1, PSMG4, RPS12, RPS18, HHEX, ETV2, GALNT11, WWOX, CDK11B, PSMA5, BZW2, IFT74, LEO1, DDT, USP51, D17WSU92E, TFAM, GCDH, CITED2, D3ERTD751E, POLR2A, ECHDC2, PRPS1, FABP5, MYG1, and ETV2, also demonstrated distinct expression patterns, as depicted in the violin plots, which consistently showed higher expression levels in the VEGFB-secreting group (Fig.**??**F). GO analysis via DAVID illuminated the top biological processes these genes are implicated in, with the Wnt signaling pathway, the determination of left/right symmetry, and cytoplasmic translation leading the processes list (Fig.**??**G). Such processes are integral to cellular communication and embryonic development, elucidating a complex network of interactions. The extensive research on the Wnt signaling pathway’s influence on Endothelium development in mouse embryos further substantiates its pivotal role in the morphogenesis and functional specialization of the vascular system, essential for CNS vasculature formation and the blood-brain barrier’s integrity^29,30^.

### Explore the communication pattern in Osteoarthritis using CLARA

The intricate interplay between inflammation and acidification in the pathogenesis of osteoarthritis (OA) has emerged as a critical area of research, illuminating how these processes synergistically contribute to the disease’s progression and severity. Studies have increasingly shown that the acidic microenvironment within OA-affected joints not only fuels the inflammatory response but also directly impacts chondrocyte metabolism and survival, thereby exacerbating cartilage degradation. Kittl et. al. demonstrates that extracellular acidification can elicit an acid-sensitive outwardly rectifying chloride current in chondrocytes. This finding is critical because cell swelling and impaired cell volume regulation under acidic conditions prevalent in OA can detrimentally affect chondrocyte viability and function, potentially accelerating disease progression^31^.Furthermore, a comprehensive analysis of articular chondrocytes from OA joints has identified a metabolic shift in chondrocytes in response to pro-inflammatory stimuli, a feature specific to OA chondrocytes and not found in non-OA chondrocytes. This metabolic alteration underlines the sensitivity of OA chondrocytes to inflammatory environments, which may be exacerbated by acidification^32^. Additionally, the pathogenic role of pro-inflammatory mediators in OA has been well established, with various cytokines such as IL1, IL6, IL17, and TNF-*α* playing crucial roles in the acute inflammatory phase of the disease. The imbalance between pro-inflammatory and anti-inflammatory cytokines characterizes this phase, suggesting that acidification could influence these inflammatory processes, further impacting chondrocyte function and disease progression^33^. Meanwhile, Dickkopf-1 (DKK1) is intricately linked to inflammation, showcasing its multifaceted role in both promoting and regulating inflammatory responses in various cellular environments. DKK1, a known antagonist of the Wnt/*β* -catenin signaling pathway, impacts inflammation and tissue repair by modulating the canonical and noncanonical Wnt pathways, which are crucial for cell proliferation, differentiation, and migration. Specifically, DKK1 competes with Wnt3a for binding to the LRP5/6 co-receptor, potentially inhibiting Wnt-mediated tissue homeostasis and affecting the balance of cell fate decisions^34^. Despite extensive research on acidification, the patterns of cell communication during OA, especially how acidification is mediated through these pathways, remain less understood. With CLARA, we could elucidate the communication mechanisms through IL6→IL6ST and DKK1→LRP6 during OA, shedding light on how these interactions influence acidification in the disease context.

With CLARA, the communication of IL6→IL6ST are mostly identified at the deep zone and middle zone in the cartilage as shown in Fig.**??** A. While the communication cells in normal have similar pattern OA and normal in middle zone in strength and count wise, there are more communication counts and high communication score in the OA than in the normal at deep zone. To further validate the communication, we group the dataset in two manners including IL6 senders vs non IL6 senders. The t-test are performed for these two groups to find the biomarkers Fig.**??** B. The GO biological process analysis is performed for top 20 biomarkers for each group Fig.**??** C. In the IL6 sender group, positive regulation of transcription from RNA polymerase II promoter, inflammatory response to wounding and cellular response to oxidative stress are highlighted in the results. As discussed by Kasre et.al., Immediate mediators of the inflammatory response are poised for gene activation through RNA polymerase II stalling^35^. And stated by Ramos-González et.al, both inflammation and oxidative stress are interrelated since one could promote the other, leading to a toxic feedback system, which contributes to the inflammatory and demyelination process^36^. In the IL6 receiver group, multiple acidifications related biological processes are highlighted in the analysis, including: Golgi lumen acidification, endosomal lumen acidification, intracellular pH reduction, vacuolar acidification and lysosomal lumen acidification. Research has shown that acidification can influence inflammation in several ways including 1. Activation of Immune Cells: Acidic environments can directly activate immune cells. Certain types of immune cells can be activated or modulated by changes in pH levels. This activation can enhance their ability to respond to pathogens but can also contribute to inflammatory processes^37^. 2. Effect on Inflammatory Mediators: Acidification can affect the production and action of various inflammatory mediators. For instance, lower pH levels can influence the production of cytokines and chemokines, molecules that play critical roles in orchestrating the immune response and are pivotal in the development and sustainment of inflammation^38^. As the IL6 is highly related with inflammation, we demonstrate how IL6 bind with IL6ST and activate the acidification in human cartilage deep zone using CLARA.

Another communication pattern, DKK1→LRP6, also displayed a similar pattern with the previous one. As they happens more frequently in OA than normal at human cartilage deep zone. However, the communication in the normal dataset has more strength than them in OA in both deep and middle zone. The GO analysis of the bio-marker genes in the sub-grouping also demonstrate the correctness of the communication cell pairs we identified. In the DKK1 sender cells, multiple wnt related biological process are highlighted including: axonogensis, canconical wnt signalling pathway and negative regulation of wnt signalling pathway etc,. The DKK1 receiver cell groups also highlight the wnt related process including: canonical wnt signaling pathway and skeletal muscle cell and fat cell differentiation. Wnt signaling, bifurcating into canonical and non-canonical pathways, regulates gene expression by modulating the stability and localization of beta-catenin or activating beta-catenin-independent mechanisms, respectively. DKK1, a potent inhibitor of this pathway, particularly affects bone metabolism by suppressing osteoblast-mediated bone formation through its interaction with LRP5/6 co-receptors, thereby inhibiting Wnt’s ability to activate its signaling cascade. This complex interplay not only underscores the significance of the Wnt/DKK1 axis in maintaining cellular and tissue equilibrium but also highlights its potential as a therapeutic target in conditions marked by dysregulated Wnt activity. This analysis demonstrates that the communication pattern between DKK1 and LRP6 is more frequent in osteoarthritis-affected cartilage than in normal cartilage, and the following GO analysis demonstrate about the correctness of our inferred cell communication using CLARA.

## Discussion

This study introduced CLARA, an innovative computational framework designed to enhance our understanding of cell-cell communication through the use of a ligand-receptor attention mechanism. By leveraging a refined self-attention model, our framework has provided detailed insights into the dynamic interplay between cells, particularly within complex biological systems such as vascular development in mouse embryos and disease progression in human osteoarthritis.

The application of CLARA to diverse datasets, including mouse embryo SeqFISH and human osteoarthritic cartilage, revealed critical communication patterns that contribute significantly to both normal physiological processes and pathological conditions. For instance, our analysis of the mouse embryo dataset highlighted how VEGF-mediated signaling pathways orchestrate vascular development, emphasizing the nuanced roles these interactions play in tissue morphogenesis and stability. In contrast, the human cartilage dataset offered a stark depiction of the communication breakdowns that may contribute to the pathophysiology of osteoarthritis, suggesting new avenues for therapeutic intervention.

One of the principal advantages of CLARA over existing tools like CellChat and stLearn is its superior specificity in identifying true signaling interactions, thus reducing the prevalence of false positives that often plague similar computational approaches. This enhanced accuracy is vital for constructing dependable models of cellular communication, which are essential for both fundamental biological research and clinical applications.

Furthermore, the scalability and adaptability of CLARA enable its application across various other biological settings, potentially providing a universal tool for exploring cell-cell communication in any tissue or disease state. Future research will aim to expand the capabilities of CLARA by incorporating multi-omic data integration, allowing for a more holistic view of cellular interactions. This could include the analysis of proteomic and metabolomic data alongside transcriptomics to offer a multi-layered understanding of cellular functions.

Additionally, longitudinal studies employing CLARA could elucidate the temporal dynamics of cell communication, shedding light on how cellular networks evolve in response to developmental signals or during disease progression. Such studies would be instrumental in deciphering the temporal aspects of cellular dialogues and could lead to the development of time-sensitive therapeutic strategies that target specific stages of disease or development.

In conclusion, our study establishes the CLARA framework as a cutting-edge tool for deciphering the complex language of cellular communication. The insights gained from this work not only advance our fundamental understanding of cellular interactions but also open up promising new directions for research and therapy in various biomedical fields.

## Methods

### Curate ligand-receptor-Transcription factor triplets

Inter-cellular responses are initiated with the binding of a ligand to its corresponding receptor, activating transcription factor in specific cell signaling pathways. The foundation for inferring Cell-Cell Communication lies in constructing a database of ligand-receptor-downstream transcription factors (TFs) triplets. In CLARA, creating the triplet database is compromised by two parts:1) Imports ligand and receptor pairs from CellChat and CellPhoneDB, and 2) Extracts receptor and corresponding TFs from the KEGG and Reactome pathways. The pathway driven receptor and downstream TFs pairs are constructed by utilizing a random walk algorithm which begins at the receptor and ends at any encountered TF of the pathway. This data collection process produces a database containing 39272 ligand-receptor-TFs triplets with 1141 unique ligand-receptor pairs for CLARA.

### Dataset pre-processing

There are two main steps for dataset pre-processing in CLARA before the cell communication inference: gene expression pre-processing and the creation of “cell sentences.”

#### Gene expression pre-processing

Initially, gene expression pre-processing refines the raw gene expression data through filtering, normalization, and a thorough quality control procedure using Scanpy^39^. These pre-processing steps aim to address gene expression dropouts and reduce the impact of extreme values produced during the wet experiments. To counter the high dropout rate characteristic of gene expression data and to minimize the effects of outliers, we apply a logarithmic transformation to the gene expression values. Specifically, the log-transformed expression value of gene *g* in cell 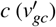 is computed after normalizing the original expression value (*v*_*gc*_) with the gene-wise median of non-zero expressions (med(*V*_*g*_ = *{v*_*g*1_, …, *v*_*gc*_|*v*_*gc*_ *>* 0*}*)) using Eq.1.

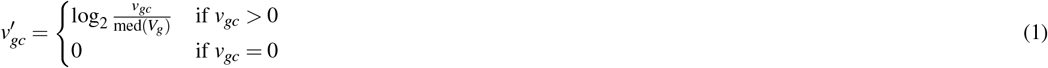

#### Cell sentence

A “cell sentence” (*cellS*_*i*_) is a sequence of vectors represented by the gene expression of a target cell (*c*_*i*_) and its neighboring cells, used to quantify cell communication from neighboring cells to the target cell. In CLARA, two cells are known as neighbor cell to each other if the distance between two cells are smaller than cell communication radius (*R*). Creating cell sentences for a dataset begins with utilizing cell locations to construct a node graph *G*(*N*) (Fig. **??** A). For a cell *c*_*i*_ in the dataset with 2-D coordinates (*x*_*i*_, *y*_*i*_), a node *n*_*i*_ is placed at (*x*_*i*_, *y*_*i*_) within the graph structure *G*(*N*). Once all cells are included, the Euclidean distances *D* = {*d*_12_, *d*_13_, …, *d*_*i j*_} between each pair of nodes (*i* and *j*) are calculated. CLARA introduces a hyper-parameter *P* to adjust the weight of the maximum communication distance when determining the final *R* (Eq. 2). The determination of *P* can either be predefined or determined by CLARA which will be explained in the following section. All cell sentences are created using two for loops as described in Algorithm 1. In the “cell sentence”, the vector for the target cell is followed by the vectors of neighboring cells, forming a sequence that enables the calculation of ligand-receptor attention. As CLARA focuses on the attention from the target cell to the neighbor cells, the order of the neighbor cells can be random.

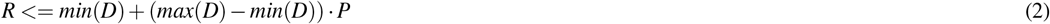

#### The choice of cell communication radius

The performance of subsequent models can be impacted by the length of the sequence. Theoretically, increasing the sequence length or the cell communication radius *R* can include more information within a single sequence, which will lead to an increase in model performance. The model performance will converge to a certain extent as the model obtains enough information within a single sequence. However, the computational load can grow exponentially with the increase in the cell communication radius. CLARA performs a grid search within a certain range of *P* to find the ‘elbow point’ between *P* and model performance as the optimal radius. (Fig. **??**).

##### Algorithm 1

Creation of cell sentences

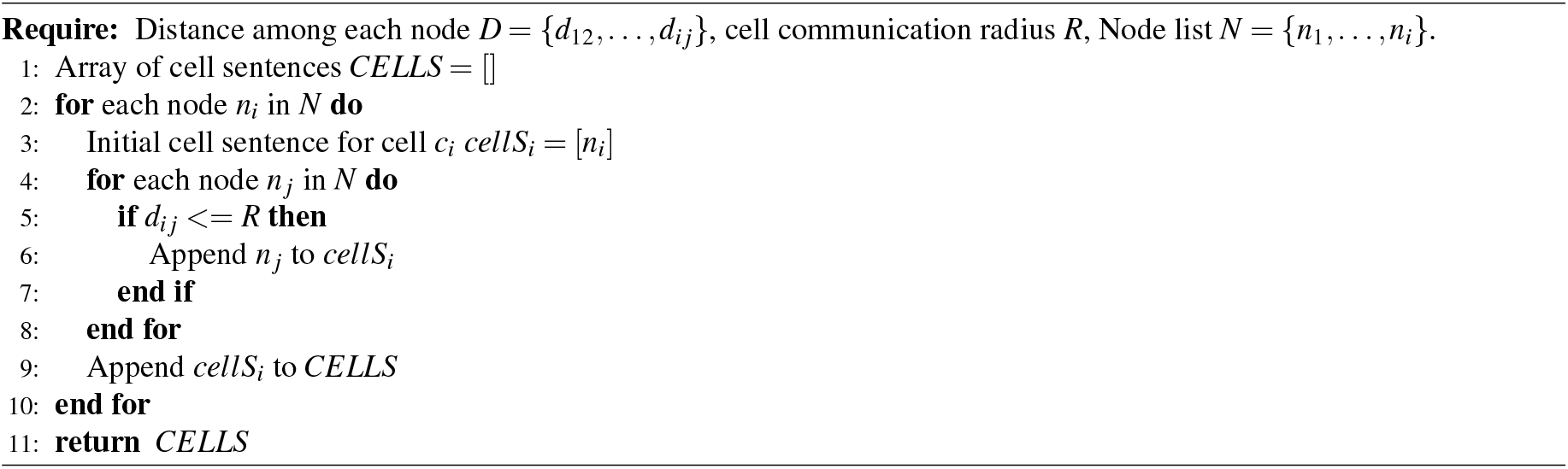

### Ligand-receptor attention mechanism

The idea of ligand-receptor attention is borrowed from the self-attention mechanism. The mechanism allows a model to weigh the importance or the impacts between different words in a sentence. This can also be applied in cell sentence to weight the how the neighbor cells impact the target cell. The theory of self-attention can be shown in Fig. **??**B with several key steps including: Query, Key, and Value, Attention Score and Weighted Sum.

#### Query, Key, and Value Vectors

For *i*th word in a sentence at *l*th layer is represented as a vector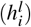, and for each word in a sentence, three different vectors are computed using separate weight matrices. These vectors are known as the Query (*Q* = *HW*^*Q*^), Key(*K* = *HW*^*K*^), and Value(*V* = *HW*^*V*^)vectors. The weight matrices are learned during the training process. Where *H* = *h*_1_, *h*_2_, …, *h*_*n*_ is the matrix of input vectors, with each row representing a word. *W*^*Q*^, *W*^*K*^ and *W*^*V*^ are the weight matrices for queries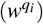, keys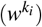 and values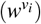, respectively. These matrices are learned during training.

#### Attention Score

The attention scores are calculated by taking the dot product of the query vector with all key vectors and scaling the result by the square root of the dimension of the key vectors (*d*_*k*_) to achieve more stable gradients. This process determines the influence or “attention” each word should have on the others. The attention score *α*_*i, j*_ quantifies the relevance between the *i*th and *j*th words, formulated as Eq.3. This ensure the scores across all words sum up to 1, Softmax function are used to turn them into probabilities that represent the importance of each word’s impact on the others (Eq.4).

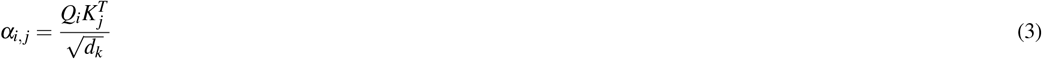

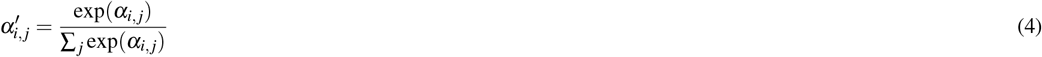

#### Weighted Sum

The normalized attention scores are used to create a weighted sum of the Value vectors as described in Eq.5. This step aggregates the information from all other words in the sentence, weighted by their relevance to the current word. The result is a new vector 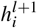 for each word that captures contextual information from the entire sentence.

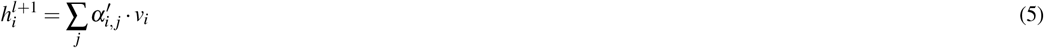

#### Ligand and receptor attention

Self-attention’s strength lies in its ability to weight how parts of the input sentence impact each other. However, the self-attention cannot be applied directly to reflect the communication through ligand and receptor pairs. As in self-attention the relationship of elements between *q*_*i*_ and *k* _*j*_ is 1:1 (Fig. **??**B), while it is a multi-valent interaction in the ligand receptor relationship. In CLARA, the self-attention is revised based on ligand and receptor mapping relationship and it is reflected by a newly added mask matrix (*W*^*l*^*r*) (Fig. **??**C). CLARA, first, linear transform the receptor of target cell vectors into Query vectors (*Q* = *H*_*r*_*W*^*Q*^), the ligand expressions for all cells in the cell sentence are transformed into a vector with same length as receptor through a trainable ligand-receptor pair mask matrix *W*^*l*^*r* by element multiplication and then through linear transformation to Key vectors *K* = *H*_*l*_*W*^*lr*^*W*^*K*^. *W*^*lr*^ is the mask matrix which is used to map relationship between ligand and receptor according to the binding priori knowledge. The Eq.6 is used to build *W*^*lr*^ while *w*_*lr*_ is a trainable parameter which will be adjusted during the training process through back prorogation. CLARA finally will follow the process self-attention to calculated the normalized attention score 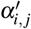 which is also the communication coefficient of communication from neighbor cell *j* to target cell *i*.

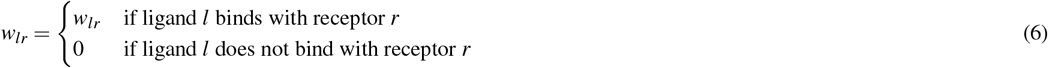

### Supervised learning for ligand-receptor attention

CLARA employs a ligand-receptor attention mechanism to determine the relevance between cells in the model, catering to two kinds of tasks: classification and regression. As shown in Fig.**??**, the model has two main components: the communication attention part and the fully connected part. The communication attention part contains one ligand-receptor attention layer and uses the ligand and receptor data, estimate the likelihood of interaction and simulate the ligand and receptor expression for target cell after communication with its neighbors. While multiple attention layers could lead to slight fluctuations in classification accuracy (1), it will also diminish the explain-ability of the model. As the revised ligand-receptor expression in the target cell could be seen as the result of communication simulated by one attention layer, the addition of more layers may complicate the understanding of which layers are most significant in driving these changes. Besides the attention part, the fully connected part is used to get additional TF information in cell gene expression. Once the model identifies the interaction patterns between ligands and receptors, it combines the outputs from both components and inputs them into another network for further processing. Two types of data can be gather to acquire the interaction between cells within cell sentences including categorical, like identifying cell types, or continuous, such as Zonal score or Biological Process (BP) scores^40^. The Adam optimizer^41^ is used to optimize the weights in the model. To prevent the over-fitting, an early stop mechanism is introduced to the training process when the validation loss starts increasing and the validation accuracy starts decreasing.

**Table 1.** Table for model classification performance compare one ligand-receptor attention layer with multiple layers using simuliated dataset and mouse embryo

| Dataset | One layer | Three layers | Five layers |
| --- | --- | --- | --- |
| Simulated dataset | 99.35% | 98.47% | 98.97% |
| Mouse embryo E1 | 84.35% | 83.45% | 81.35% |
| Mouse embryo E2 | 87.31% | 79.35% | 81.25% |
| Mouse embryo E3 | 82.35% | 76.65% | 80.32% |

### The inference of cell-cell communication

After the training stops, we infer the communication score with a heuristic rule by taking the communication coefficient *coe f*, the ligand and receptor expression value, and corresponding downstream TFs expression in each cell pairs. Intuitively, for effective communication between sender and receiver cells via ligands and receptors, the expression levels of a ligand in the sender cell and a receptor in the receiver cell should be high. Conversely, the expression levels of a receptor in the sender cell and a ligand in the receiver cell should be low. In the heuristic rule, two aspect are employed to further refine the inferred cell communication. The first one is adding the expression relationship pattern between ligand and receptor in sender cell and receiver cell to the communication score. CLARA compute the communication score from *c* _*j*_ to *c*_*i*_ through ligand and receptor pair *lr* using Eq.7 where ligand and receptor pair expression value in both cells *i* and *j* are denoted by (*l*_*i*_, *r*_*i*_) and (*l* _*j*_, *r* _*j*_), respectively. The *LRE*(·) shown in Eq.8 is the function computing the ligand and receptor communication strength where the 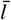 and 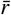 is the average expression of ligand and receptor. To prevent all the values from becoming close to zero and reduce the effect of small value that will cause the result explosion, we use Laplace Smoothing with the average of ligand and receptor expression. One more refining method filters communication by only confirming it when the downstream is activated. CLARA utilizes two filtering strategies to minimize false positives in this method. The first strategy requires that at least one downstream transcription factor (TF) be expressed in the receiving cell *c*_*i*_. This is followed by comparing the ligand and receptor expression levels to the average expression of cells in the set *N* (*N* = *c*_1_, *c*_2_, …, *c*_*n*_). These rules are outlined in Eq.9 where *TF*_*ri*_ represents the expression of all transcription factors downstream of the receptor in cell *c*_*i*_. The final communication score is then calculated using Eq.10

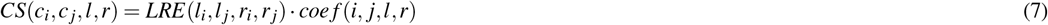

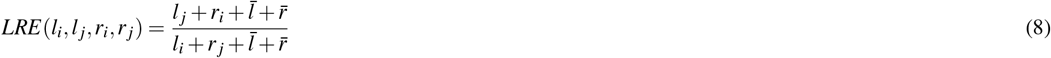

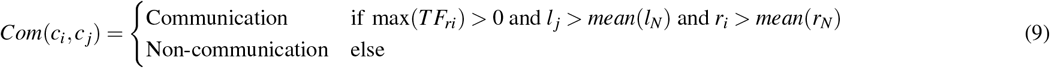

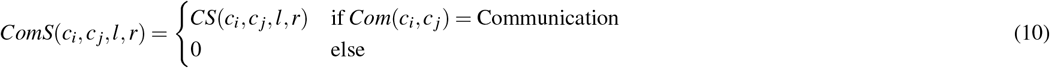

### The inference of cell type communication

For the scenario that cell type information is provided, to generalize specific cell-cell communications to cell group-level interactions, CLARA aggregate all the communication scores from group A to group B by grouping feasible ligand and receptor pairs. We employ a permutation test to determine the statistical significance of the inferred cell group communication. The permutation test involves random permutations of cell labels followed by the recalculation of the average group communication score between A and B via the communication ligand l and receptor pair r Eq.11. Here, *CS*(*A, B, l, r, d*_*i*_) represents the mean of communication score for cells in group *α* communicating with cells in group *β* through a particular ligand and receptor pair Eq.12. *CS*^*′*^(*A, B, l, r*) corresponds to the mean of communication score after the cell labels have been randomly permuted. We repeat this process for a total of Q random samples, with Q default set to 100 and the default p-value threshold for the statistically significant communication is set to 0.05.

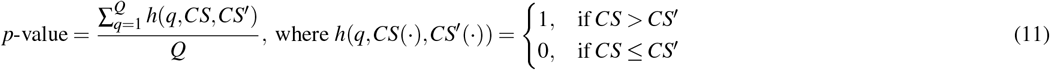

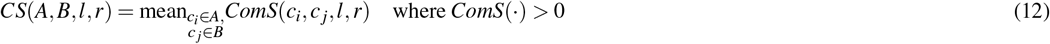

### Human cartilage zonal score

Zonal score is a kind of score that uses to hypothetically inferred the cell location from the surface of the human cartilage to the deep zones near the bone. Multiple studies have state that human cartilage can be split into three zones including Superficial zone (S), Middle Zone (M) and Deep Zone (Z) based on the expression pattern of multiple marker genes. A list of zonal marker genes in the three major zones of the human cartilage is provided in Table2. The Zonal Scores is calculated by 13

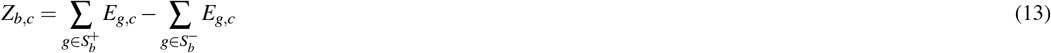

where *E*_*g,c*_ is the expression value of the gene *g* of the cell *c*, 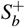 and 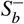 represent the positive regulation and negative regulation genes annotated for the biology defining the zonal index *b* respectively^42^. The higher score stands for the cell more approaching to the surface.

## Code availability

CLARA is publicly available as a Python package. Source codes, as well as tutorials have been deposited at the GitHub repository (https://github.com/Harry-Wang12/CLARA/).

**Table 2.** Table for zonal marker genes in human cartilage

| Superficial Marker ($S_b^+$) | Deep Marker ($S_b^-$) |
| --- | --- |
| EFEMP1 | CLEC3A |
| IGFBP5 | SLC44A2 |
| CHI3L1 | GLT25D2 |
| THBS4 | SLC7A5 |
| MYO1B | LECT1 |
| BMPER | F13A1 |
| PDE3B | VAV3 |
| OGN | COL10A1 |
| PALMD | IBSP |
| FGFR2 | SPP1 |
| FGL2 |  |
| TNC |  |
| PRG4 |  |

## Acknowledgements (not compulsory)

Acknowledgements should be brief, and should not include thanks to anonymous referees and editors, or effusive comments. Grant or contribution numbers may be acknowledged.

## Author contributions statement

Must include all authors, identified by initials, for example: A.A. conceived the experiment(s), A.A. and B.A. conducted the experiment(s), C.A. and D.A. analysed the results. All authors reviewed the manuscript.

## Additional information

